# The Mba1 homologue of *Trypanosoma brucei* is involved in the biogenesis of oxidative phosphorylation complexes

**DOI:** 10.1101/2022.12.21.521360

**Authors:** Christoph Wenger, Anke Harsman, Moritz Niemann, Silke Oeljeklaus, Corinne von Känel, Salvatore Calderaro, Bettina Warscheid, André Schneider

## Abstract

Consistent with other eukaryotes, the *Trypanosoma brucei* mitochondrial genome encodes mainly hydrophobic core subunits of the oxidative phosphorylation system. These proteins must be co-translationally inserted into the inner mitochondrial membrane and are synthesized by the highly divergent trypanosomal mitoribosomes, which have a much higher protein to RNA ratio than any other ribosome. Here, we show that the trypanosomal ortholog of the mitoribosome receptor Mba1 (TbMba1) is essential for normal growth of procyclic trypanosomes but redundant in the bloodstream form, which lacks an oxidative phosphorylation system. Proteomic analyses of TbMba1-depleted mitochondria from procyclic cells revealed reduced levels of many components of the oxidative phosphorylation system, most of which belong to the cytochrome c oxidase (Cox) complex, three subunits of which are mitochondrially encoded. However, the integrity of the mitoribosome and its interaction with the inner membrane were not affected. Pulldown experiments showed that TbMba1 forms a dynamic interaction network that includes the trypanosomal Mdm38/Letm1 ortholog and a trypanosome-specific factor that stabilizes the CoxI and CoxII mRNAs. In summary, our study suggests that the function of Mba1 in the biogenesis of membrane subunits of OXPHOS complexes is conserved among yeast, mammalian, and trypanosomes, which belong to two eukaryotic supergroups.

## Introduction

Mitochondria or mitochondria-derived organelles are a defining feature of eukaryotic cells. They originated from a single endosymbiotic event between an α-proteobacterium and an archaeal host cell, 1.7 to 2.1 billion years ago^1–5^. During the endosymbiont-to-mitochondria transition, most of the endosymbiont’s genes were either lost or transferred to the genome of the host cell^1,6^. However, a small number of genes were retained in the mitochondrial DNA (mtDNA). Thus, mitochondria contain a complete transcription and translation machinery, including ribosomes termed mitoribosomes.

Many of the genes that were retained in the mtDNA are highly hydrophobic core subunits of the oxidative phosphorylation (OXPHOS) complexes^1^. Thus, the mtDNA of *Saccharomyces cerevisiae* encodes for eight proteins, seven of which are hydrophobic components of the OXPHOS complexes^7–9^. These proteins are co-translationally inserted into the inner mitochondrial membrane (IM) by the protein oxidase assembly 1 (Oxa1)^10^, a homolog of the bacterial insertase YidC^11,12^. Oxa1 has a large hydrophilic, matrix-facing C-terminal domain that was shown to be in close proximity to the mitoribosomal proteins (MRPs) uL23 and uL24 of the mitoribosomal large subunit (mt-LSU)^13,14^. This is in line with the fact that in *S. cerevisiae* all mitoribosomes are associated with the IM^10^.

Interestingly, mitoribosomes remain associated with the IM even in the absence of Oxa1^15^, indicating that additional factors are involved in the IM association of mitoribosomes. Indeed, several such factors have been discovered. The first one identified, termed multi-copy bypass of AFG3 or (Mba1)^16,17^, is peripherally associated with the IM on the matrix side and serves as a mitoribosome receptor^16^. It is in close proximity to the MRPs uL22 and uL29^13,14^ and aligns the peptide exit tunnel of the mitoribosome with the insertion site of Oxa1 at the IM^13,18^. Mba1 interacts with the mitochondrial distribution and morphology protein 38^19^ (Mdm38, in other organisms referred to as Letm1), an integral IM protein also involved in the association of mitoribosomes with the IM^20^. Mba1 and Mdm38 might additionally be involved in the recruitment of mRNA stabilization and translation factors^19^. Finally, the expansion segment 96-ES1 of the yeast mitochondrial 21S rRNA has also been shown to contribute to the IM association of mitoribosomes^21^. An impaired co-translational IM integration leads to decreased activity of the OXPHOS complexes, since they rely on mitochondrially encoded subunits^15,16,22^. Mdm38 was additionally shown to have a second role in the cell’s K^+^/H^+^ homeostasis^23^.

Mitochondrial protein biogenesis, including co-translational IM integration of mitochondrially encoded proteins, is best understood in *S. cerevisiae* and mammals, both belonging to the eukaryotic supergroup Amorphea^24^. The arguably next best understood system concerning mitochondrial protein biogenesis, is the unicellular parasite *Trypanosoma brucei*. It belongs to a different eukaryotic supergroup, called the Discoba, which is essentially unrelated to the Amorphea^24^. Indeed, the work on mitochondrial protein biogenesis in *T. brucei* showed that, even though the general pathways are conserved, the involved complexes are astonishingly diverse^25–28^.

*T. brucei* undergoes a complex life cycle, during which it adapts to very different environments. In the midgut of its insect vector, the Tsetse fly, the procyclic form (PCF) of *T. brucei* encounters glucose-poor and amino acid-rich conditions^29^. Here, it uses proline as its main energy source^30,31^. The OXPHOS system is fully functional and the single mitochondrion of *T. brucei* forms an intricate network spanning the whole cell^32,33^. As in *S. cerevisiae*, most of the genes encoded on the trypanosomal mtDNA, termed kinetoplast DNA (kDNA), are subunits of the OXPHOS complexes. 12 of the 18 protein-coding transcripts encode for subunits of the OXPHOS complexes^34–36^.

On the other hand, the bloodstream form (BSF) of *T. brucei*, found in the blood of its mammalian host, produces energy by glycolysis using glucose as an energy source^32,33^. Here, the mitochondrion is heavily reduced and OXPHOS complexes are absent^33,37^, with the exception of the ATP synthase, which however functions in reverse in an ATP-consuming manner to maintain the membrane potential^38,39^.

To date, one of the least understood processes of mitochondrial protein biogenesis in *T. brucei* is the integreation of mitochondrially encoded proteins into the IM. Here, we identify the protein Tb927.7.4620 as the trypanosomal ortholog of Mba1 (TbMba1). We present data showing that TbMba1 is essential for the maturation and/or stability of trypanosomal OXPHOS complexes. Furthermore, we show that it interacts with the trypanosomal ortholog of Letm1/Mdm38 and with the trypanosome-specific Tb927.11.16250, a stabilizing factor of the mitochondrial mRNAs encoding cytochrome c oxidase subunit I (CoxI) and CoxII.

## Results

### Tb927.7.4620 is an ortholog of yeast Mba1

A recent publication identified the *Leishmania major* protein Lmjf1.14.0080 as the ortholog of yeast Mba1^40^. *L. major*, like *T. brucei*, is a trypanosomatid^41^ and thus belongs to the Discoba^24^. When we did BLAST searches for the ortholog of Lmjf.14.0080 in *T. brucei*, we identified Tb927.7.4620 as its homologue. Excluding the predicted N-terminal presequences (MitoProt II server: https://ihg.gsf.de/ihg/mitoprot.html), the two proteins show an identity of 62.5% and a similarity of 76.0% (Fig. S1A). An alignment between Tb927.7.4620 and yeast Mba1 (Uniprot: P38300), excluding the predicted N-terminal presequences, reveals a 18.3% identity and 31.3% similarity (Fig. S1B). This strongly suggests that Tb927.7.4620 is the Mba1 ortholog in *T. brucei* which is why we termed it TbMba1.

### TbMba1 is an essential membrane-associated mitochondrial protein in PCF *T. brucei*

To functionally analyse TbMba1, a *T. brucei* PCF cell line, allowing inducible RNA interference (RNAi) against the open reading frame (ORF) of TbMba1, was established. The growth of uninduced (-Tet) and induced (+Tet) RNAi cells was analysed over six days. The knockdown of TbMba1 in PCF *T. brucei* led to a strong growth retardation starting after two days of induction, indicating that TbMba1 is essential for normal growth in this life cycle stage (Fig. 1A). In contrast, in *S. cerevisiae*, depletion of Mba1 alone does not cause a growth defect^18^. Interestingly, we were able to generate a double knockout (dKO) *T. brucei* BSF cell line, replacing the two alleles of TbMba1 with the antibiotic resistance cassettes of hygromycin and blasticidin, respectively (Fig. 1B, S2A). TbMba1 is therefore not required for normal growth of BSF *T. brucei* indicating it has a PCF specific function. This is in line with a previous study, which showed that TbMba1 expression is 11-fold lower in BSF cells when compared to the PCF of *T. brucei*^42^.

**Figure 1.**
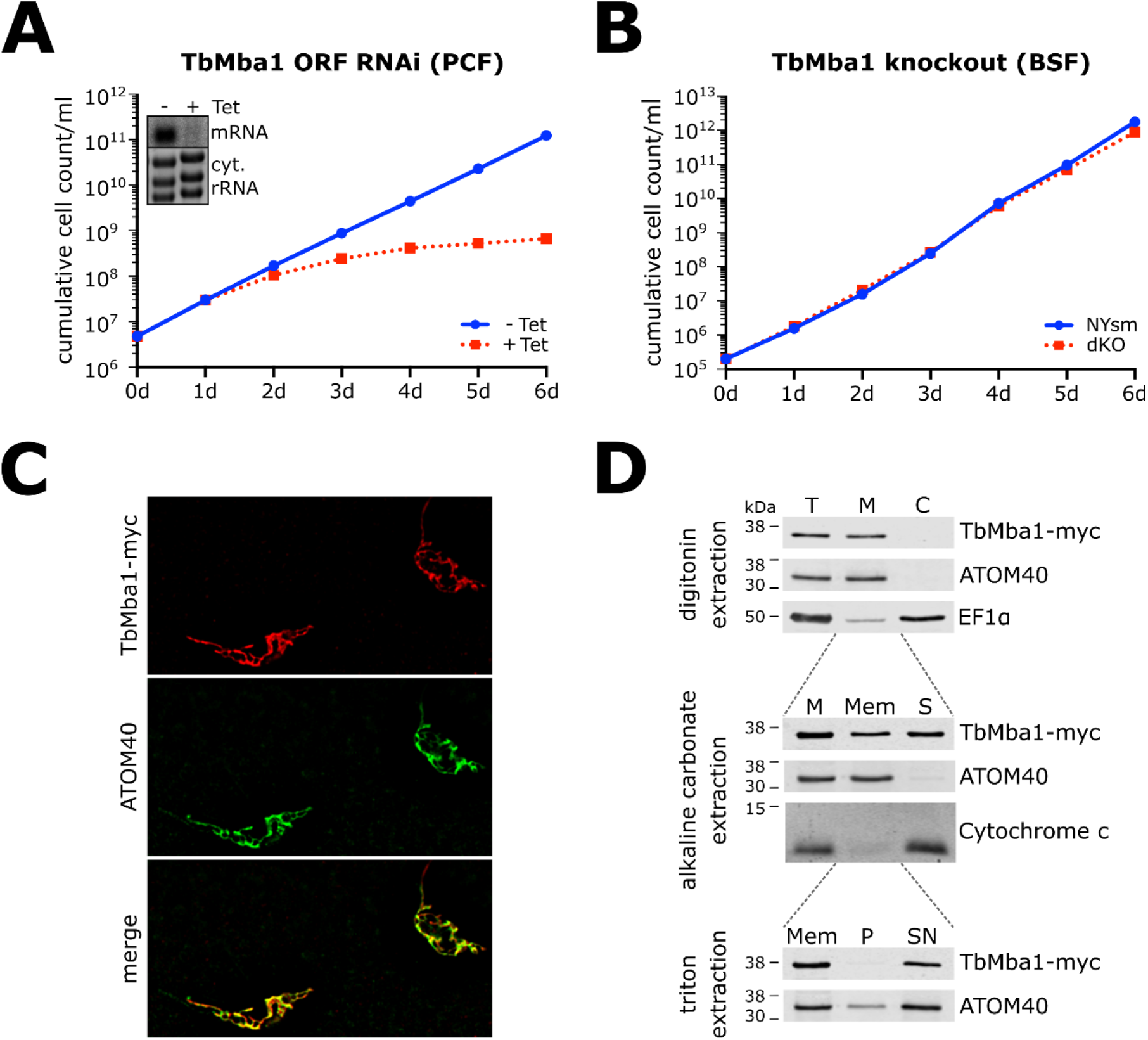
**A**: Triplicate of growth curve of uninduced (-Tet) and induced (+Tet) TbMba1 RNAi PCF cell line targeting the ORF of TbMba1. Inset shows a northern blot of total RNA from uninduced and two days induced cells, probed for the ORF of TbMba1. Ethidium bromide stained cytosolic (cyt.) rRNAs serve as loading control. **B**: Growth curve of wild type NYsm BSF cell line and TbMba1 dKO BSF cell line. **C**: IF analysis of TbMba1-myc exclusive expressor. ATOM40 serves as a mitochondrial marker. **D**: Digitonin extraction: Whole cells (T) of two days induced TbMba1-myc exclusive expressor cells were treated with 0.015% digitonin and differential centrifugation to separate a mitochondrial fraction (M) from a cytosolic fraction (C). Immunoblot analysis using anti-myc antibody to detect TbMba1-myc was performed. Anti-ATOM40 and anti-EF1α antibodies were used to detect a mitochondrial and cytosolic control, respectively. Alkaline carbonate extraction: A digitonin-extracted mitochondrial fraction was generated and solubilized in an alkaline carbonate extraction at pH 11.5. Differential centrifugation resulted in a pellet (Mem) and soluble (S) fraction that were analysed by immunoblotting. Anti-myc antibody was used to detect TbMba1-myc. Anti-ATOM40 and anti-Cytochrome c antibodies were used as controls for a membrane integral and membrane-associated protein, respectively. Triton extraction: Half of the Mem fraction from the alkaline carbonate extraction was solubilized in 1% triton. Differential centrifugation resulted in a pellet (P) and soluble (SN) fraction. Immunoblot analysis was performed using anti-myc and anti-ATOM40 antibody.

In a previous proteomics study, TbMba1 was shown to be mitochondrially localized^43^. To confirm this finding, TbMba1 was C-terminally myc-tagged (TbMba1-myc) and ectopically expressed in a PCF RNAi cell line targeting the 3’-untranslated region (3’-UTR) of the endogenous TbMba1 mRNA. This results in essentially exclusive expression of TbMba1-myc. TbMba1-myc complemented the growth phenotype caused by the depletion of the endogenous TbMba1, proving that the tag does not interfere with TbMba1 function (Fig. S2BC). Next, we analysed the localization of TbMba1-myc in the same cell line using immunofluorescence (IF) microscopy. After two days of induction, TbMba1-myc localized to the reticulated mitochondrial structure spanning the whole cell of PCF *T. brucei*. The very same structure was also stained by the mitochondrial marker atypical translocase of the outer mitochondrial membrane 40 (ATOM40) (Fig. 1C).

Furthermore, the TbMba1-myc exclusive expressor cell line was subjected to a digitonin fractionation, in which a mitochondria-enriched pellet was separated from a soluble fraction enriched in cytosolic proteins. These fractions were analysed by SDS-PAGE and immunoblotting and showed the expected co-fractionation of TbMba1-myc with ATOM40. The cytosolic marker eukaryotic translation elongation factor 1α (EF1α) however is exclusively detected in the cytosolic fraction, showing successful separation of organelles and cytosol (Fig. 1D, top panel). A fraction of the mitochondria-enriched pellet was subjected to an alkaline carbonate extraction at high pH, which allows separation of soluble proteins from membrane-integral proteins. After differential centrifugation, a large fraction of TbMba1-myc was recovered in the soluble fraction together with the peripheral membrane protein marker cytochrome c. However, a significant amount of the protein was also recovered in the membrane fraction (Mem) together with the integral membrane protein ATOM40 (Fig. 1D, middle panel). A subsequent Triton-X-100 extraction of the Mem fraction resulted in a complete solubilization of TbMba1-myc, showing that the fraction of TbMba1 present in the Mem sample was not aggregated (Fig. 1D, bottom panel). Thus, while most of TbMba1 is soluble, a smaller fraction appears to be associated with the membrane.

In summary, TbMba1 is essential for normal growth in PCF *T. brucei*, but dispensable in the BSF. C-terminally myc-tagged TbMba1 is functional and localizes to the mitochondrion, where it seems to be peripherally associated with the mitochondrial membrane. Thus, it behaves similar as Mba1 of *S. cerevisiae*, which is a peripheral IM protein that faces the matrix^16^.

### TbMba1 depletion preferentially affects OXPHOS components

In *S. cerevisiae*, Mba1 interacts with the mitoribosome aligning its exit tunnel with the IM insertase Oxa1 and mediates the integration of OXPHOS subunits into the IM^10^. However, the loss of yeast Mba1 only causes a growth defect and inactivation of OXPHOS when Oxa1 is abolished at the same time^22^. To determine the effect of TbMba1 ablation on a global level in *T. brucei*, a quantitative proteomic analysis of the steady-state levels of mitochondrial proteins in the TbMba1 ORF RNAi cell line was performed. The cell line was grown under stable isotope labelling by amino acids in cell culture (SILAC) conditions in medium containing different stable isotope-labelled forms of arginine and lysine. After three days of induction, equal cell numbers of induced and uninduced cultures, grown in presence of either light or heavy amino acids, were mixed. Mitochondria-enriched pellets were obtained by digitonin extraction and analysed by quantitative mass spectrometry (MS). All proteins that were detected in at least two of three independent biological replicates and that have previously been identified to be mitochondrial proteins^35,37,43^ are depicted in the volcano plot in Fig. 2A.

**Figure 2.**
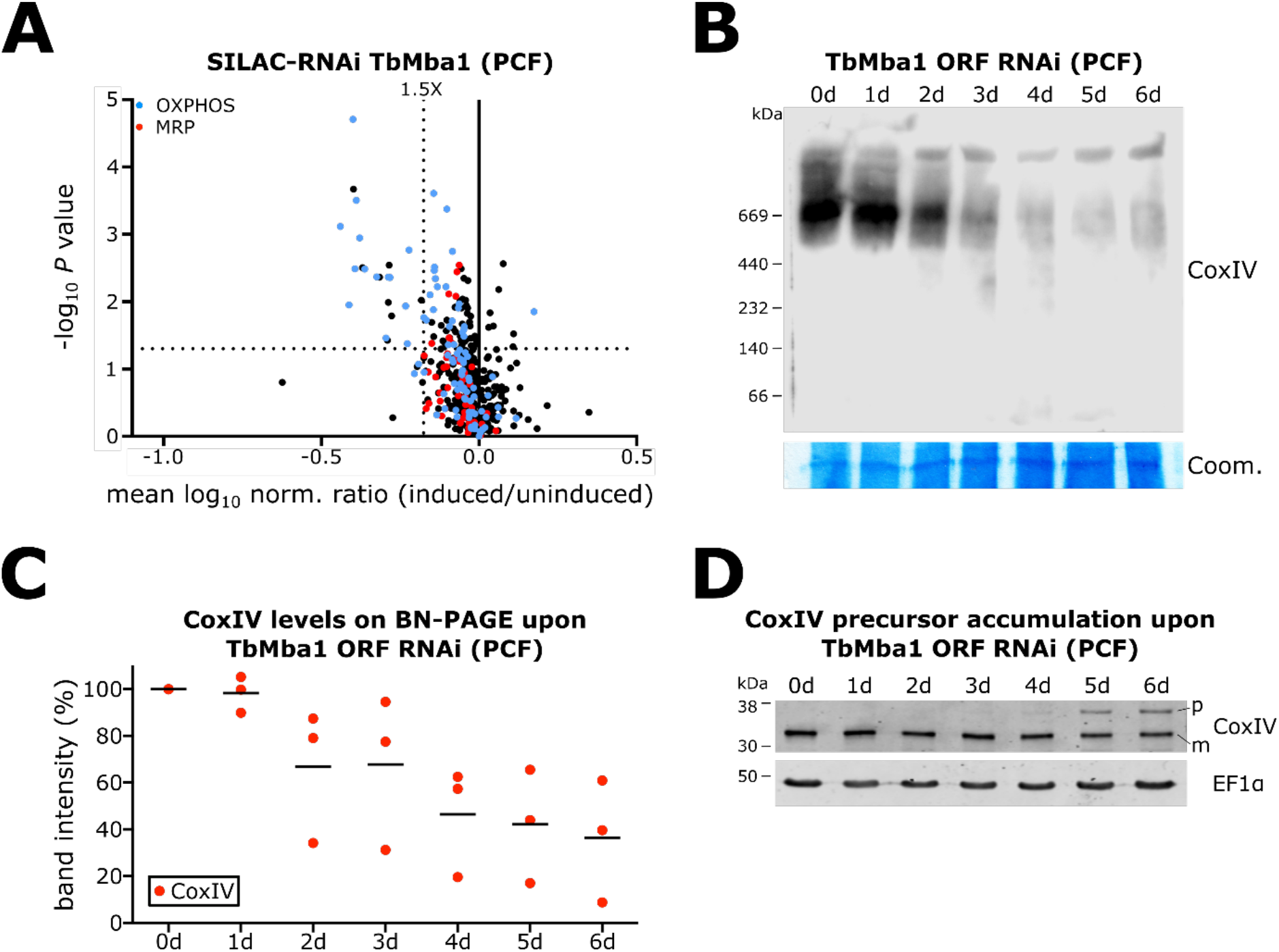
**A**: Mitochondria-enriched extracts of uninduced and three days induced TbMba1 ORF RNAi PCF cells, grown under SILAC-conditions, were mixed and analysed by quantitative MS. The volcano plot shows the mean log^10^ of the normalized (norm.) ratio (induced/uninduced) of mitochondrial proteins that were detected in at least two of three independent biological replicates, plotted against the corresponding -log^10^ *P* values (two-sided t-test). TbMba1 was only detected in one replicate and does not appear in the volcano plot. The vertical dotted line indicates 1.5-fold downregulation. The horizontal dotted line indicates a t-test significance level of 0.05. **B**: BN-PAGE and immunoblot analysis of complex IV in the TbMba1 ORF RNAi PCF cell line. Mitochondria-enriched fractions were prepared after zero to six days (0d-6d) of RNAi induction and separated on a BN-PAGE. Complex IV was detected by probing with anti-CoxIV antibody. Coomassie (Coom.) serves as loading control. **C**: Densitometric quantification of CoxIV signals as shown on BN-PAGE in B. Three independent biological replicates were measured. The levels of uninduced cells (0d) were set to 100%, the mean for each day is indicated. **D**: Whole cell extracts were harvested over zero to six days (0d-6d) of TbMba1 ORF RNAi induction and analysed by SDS-PAGE and immunoblotting, using anti-CoxIV and anti-EF1α antibodies. The precursor (p) and mature (m) forms of CoxIV are indicated and EF1α serves as a loading control.

The RNAi target, TbMba1, was downregulated 19-fold in one replicate, but not detected in the other two (Table S1), it is therefore not depicted in the volcano plot. 23 mitochondrial proteins were significantly more than 1.5-fold downregulated. 14 of them (61%) are known components of OXPHOS complexes, according to Zíková *et al*. (2017)^37^ (Fig. 2A) and 12 of the 14 OXPHOS subunits are subunits of cytochrome c oxidase (Cox), also termed complex IV. Hence, the proteins most affected by downregulation of TbMba1 are predominantly complex IV subunits. Interestingly, none of the MRPs detected in the same analysis was downregulated more than 1.5-fold, suggesting that depletion of TbMba1 does not affect the integrity of the mitoribosomes. Of the other eight mitochondrial proteins (35%) that were significantly downregulated more than 1.5-fold, one is annotated as a nucleobase/nucleoside transporter 8.2 (Tb927.11.3620), one as a putative pteridine transporter (Tb927.10.9080), one as a putative amino acid transporter (Tb927.8.8300), and five are trypanosomatid-specific proteins (Tb927.7.7090, Tb927.10.6200, Tb927.10.4240, Tb927.5.2560, Tb927.11.16510).

Downregulation of complex IV was verified by analysing solubilized mitochondria-enriched fractions of uninduced and induced TbMba1 ORF RNAi cells by blue native (BN)-PAGE and subsequent immunoblotting using an antiserum specific against CoxIV (Fig. 2B). Quantification of triplicate BN-PAGE gels (Fig. 2C) revealed a decreased signal to 67% after two days of RNAi induction. These results support that TbMba1, just as Mba1 in *S. cerevisiae*, is involved in the biogenesis of OXPHOS complexes.

To exclude that the observed loss of OXPHOS subunits is due to a possibly indirect decrease of mitochondrial protein import, whole cell samples were harvested every day over a six-day period after TbMba1 ORF RNAi induction and analysed by SDS-PAGE and immunoblotting. CoxIV contains an N-terminal presequence that is cleaved off after successful import, resulting in a shorter mature protein. Accumulation of uncleaved CoxIV precursor in the cytosol is an established hallmark of a mitochondrial protein import defect^26,44,45^. CoxIV immunoblots of whole cell extracts revealed an accumulation of CoxIV precursor after five days of TbMba1 ORF RNAi induction (Fig. 2D), three days after the onset of the growth phenotype (Fig. 1A). This means that the subunits of the OXPHOS complexes, including the ones of complex IV, are depleted well before a mitochondrial protein import defect is seen (Fig. 2ABC). Thus, TbMba1 seems to have a direct role in the biogenesis of OXPHOS subunits, but is not involved in mitochondrial protein import.

The biogenesis of OXPHOS complexes relies on the proper translation of mitochondrially encoded subunits of OXPHOS complexes by the mitoribosome. Thus, similar phenotypes as the ones observed after downregulation of TbMba1 might also be expected when we directly interfere with mitochondrial translation. To test this, we knocked down TbMRPL22, a protein of the large subunit of the mitoribosome. Fig. 3A shows that as expected for a MRP, this causes a growth retardation starting three days after induction. A SILAC-RNAi experiment with the TbMRPL22 ORF RNAi cell line, using the same procedure as explained above for the TbMba1 SILAC RNAi was done. The cells were harvested on day three after induction. All proteins that were detected in at least two of three independent biological replicates and that have previously been identified to be mitochondrial proteins^35,37,43^, are depicted in the volcano plot in Fig. 3B. TbMRPL22, the target of the RNAi, is efficiently downregulated more than tenfold (Table S2). Moreover, 52 mitochondrial proteins were significantly downregulated more than 1.5-fold. As expected, many (29) of these proteins are MRPs suggesting that, in contrast to TbMba1 ablation, depletion of TbMRPL22 affects the integrity of the mitoribosomes. Of the remaining 23 proteins, 11 are OXPHOS subunits, of which nine belong to complex IV. Thus, in regard to OXPHOS components ablation of TbMRPL22 essentially matches the effects seen after depletion of TbMba1.

**Figure 3.**
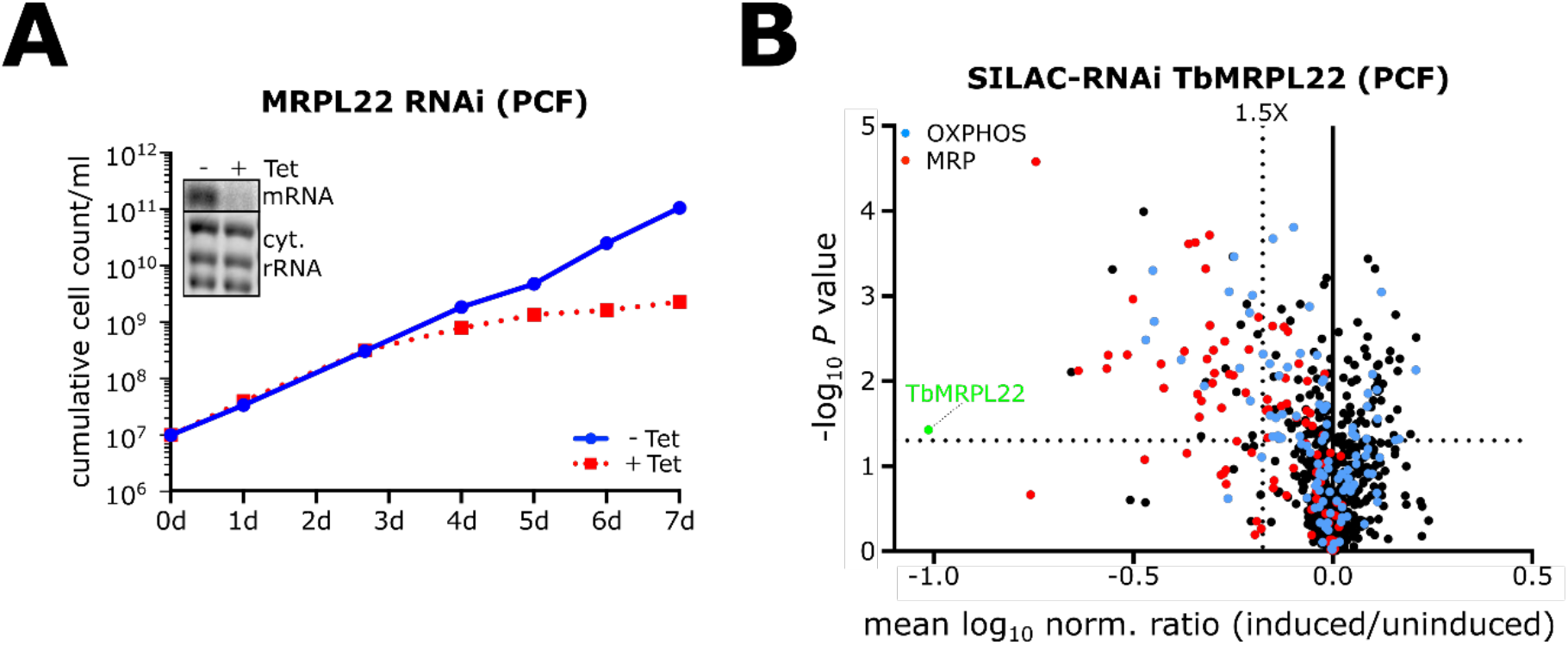
**A**: Growth curve of uninduced (-Tet) and induced (+Tet) TbMRPL22 RNAi PCF cell line targeting the ORF of TbMRPL22. Inset shows northern blot of total RNA from uninduced and two days induced cells, probed for the ORF of TbMRPL22. Ethidium bromide stained cytosolic (cyt.) rRNAs serve as loading control. **B**: Mitochondria-enriched extracts of uninduced and three days induced TbMRPL22 ORF RNAi PCF cells, grown under SILAC-conditions, were mixed and analysed by quantitative MS. The volcano plot shows the mean log^10^ of the normalized (norm.) ratio (induced/uninduced) of mitochondrial proteins that were detected in at least two of three independent biological replicates, plotted against the corresponding -log^10^ *P* values (two-sided t-test). The vertical dotted line indicates 1.5-fold downregulation. The horizontal dotted line indicates a t-test significance level of 0.05.

As discussed above, depletion of TbMba1 only marginally affects the levels of MRPs suggesting that the mitoribosomes are still intact. This allows to investigate whether they are still associated with the mitochondrial IM. To that end, we produced two TbMba1 ORF RNAi cell lines expressing in situ HA-tagged MRPs mL78 (mL78-HA) or L20 (L20-HA), respectively (Fig. 4AB). Mitochondria-enriched pellets of these two cell lines, obtained by digitonin extraction, were subjected to 15 freeze-thaw cycles followed by differential centrifugation to obtain a membrane (Mem) and a matrix (Mat) fraction. Subsequently, the fractions were analysed by SDS-PAGE and immunoblotting. The matrix protein TbmHsp70 and the IM protein TbTim17 were used as the matrix and membrane markers, respectively. Interestingly, both mL78-HA and L20-HA remained predominantly in the Mem fraction in both uninduced and up to six days induced cells. Thus, depletion of Mba1 is not sufficient to abolish the membrane-association of trypanosomal mitoribosomes (Fig. 4C).

**Figure 4.**
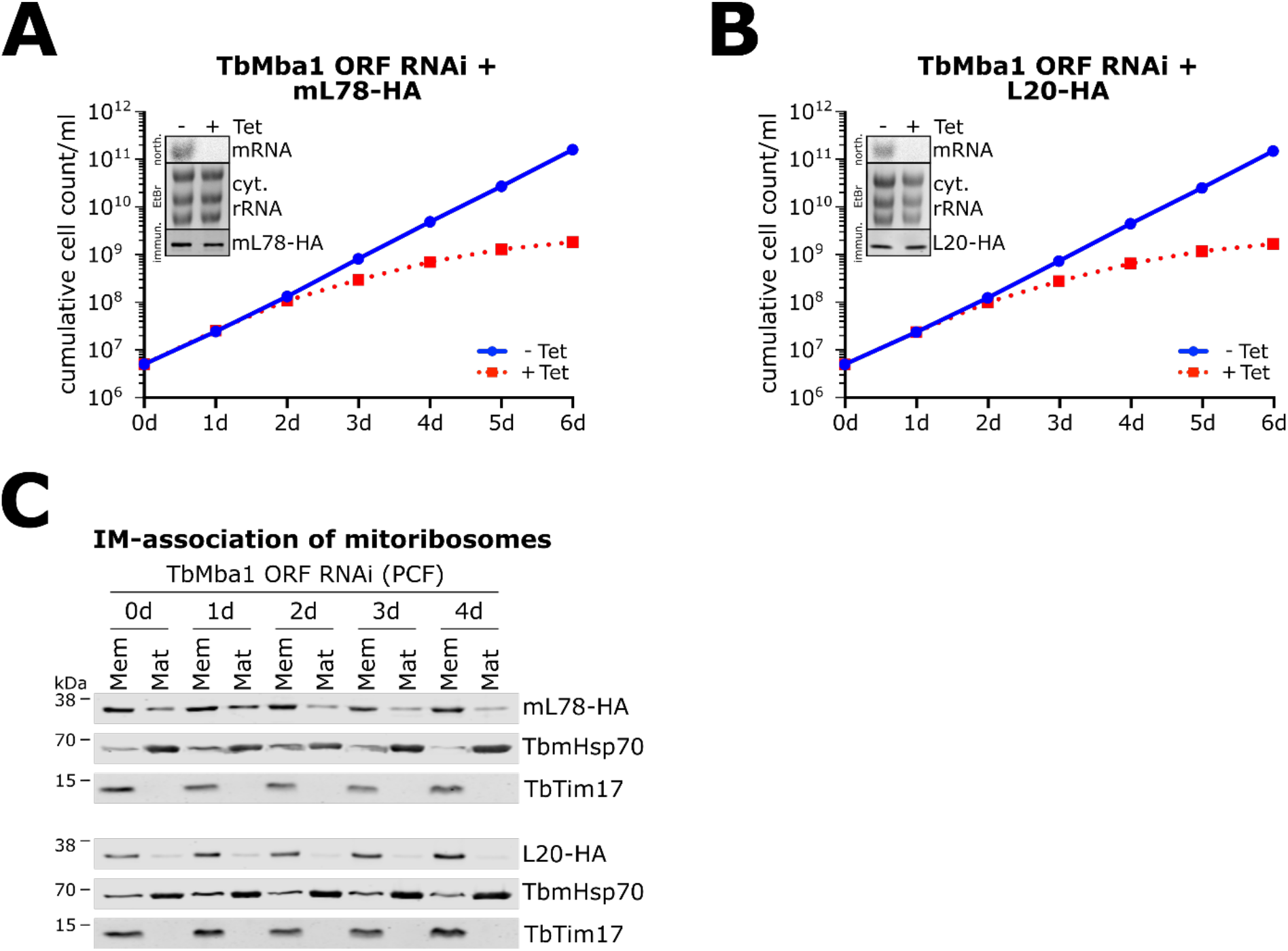
**A:** Growth curve of uninduced (-Tet) and induced (+Tet) TbMba1 RNAi PCF cell line targeting the ORF of TbMba1 in the background of mL78-HA in-situ expression. Inset shows northern blot (north.) of total RNA from uninduced and two days induced cells, probed for the ORF of TbMba1. Ethidium bromide (EtBr) stained cytosolic (cyt.) rRNAs serve as loading control. Immunoblot (immun.) analysis of whole cell extracts, probed with anti-HA antibody, to show expression of mL78-HA. **B**: As in A, but L20-HA is in-situ expressed instead of mL78-HA. **C**: Immunoblot analysis of two cell lines, either in-situ expressing mL78-HA or L20-HA in the background of TbMba1 ORF RNAi in PCF *T. brucei*. The RNAi was induced from zero to four days (0d-4d) and mitochondria-enriched extracts were separated into membrane (Mem) and matrix (Mat) fractions by freeze-thaw cycles. The tagged proteins were detected using anti-HA antibody. Detection with anti-TbmHsp70 and anti-TbTim17 served as matrix and membrane controls, respectively.

This suggests that *T. brucei* mitoribosomes use multiple factors to associate with the IM^40^ as is the case in *S. cerevisiae*^15,19^. The essential function of TbMba1 goes therefore beyond contributing to a physical connection that associates the mitoribosome to the IM.

In summary, ablation of TbMba1 in PCF *T. brucei* causes the depletion of OXPHOS components with a preference for complex IV subunits. This phenotype can be mimicked when mitochondrial translation is directly abolished by knockdown of the MRP TbMRPL22. Knockdown of TbMba1 however does neither affect steady-state levels of MRPs nor the physical connection of the mitoribosome with the IM.

### TbMba1 interacts transiently with TbLetm1 and Tb927.11.16250

To find possible interaction partners, a SILAC-based quantitative MS analysis of a co-immunoprecipitation (SILAC-CoIP) was performed using the cell line that exclusively expresses TbMba1-myc (Fig. S2C). In order to identify proteins that stably interact with TbMba1, equal cell numbers of uninduced and two days induced cultures, grown in presence of either light or heavy isotope-containing arginine and lysine, were mixed. Subsequently, mitochondria-enriched fractions were generated, solubilized, and subjected to CoIP. The eluates were analysed by quantitative MS and all mitochondrial proteins^35,37,43^ that were detected in at least two of three independent biological replicates are depicted in the volcano plot in Fig. 5A. The bait TbMba1 was enriched 98-fold demonstrating that the CoIP has worked (Table S3). However, no other protein was enriched more than fivefold indicating that TbMba1 does not efficiently form a stable complex with other proteins. This result is supported by the fact that no high molecular weight complex was detected when a solubilized, mitochondria-enriched fraction of the TbMba1-myc exclusive expressor cell line was analysed by BN-PAGE (Fig. S3).

**Figure 5.**
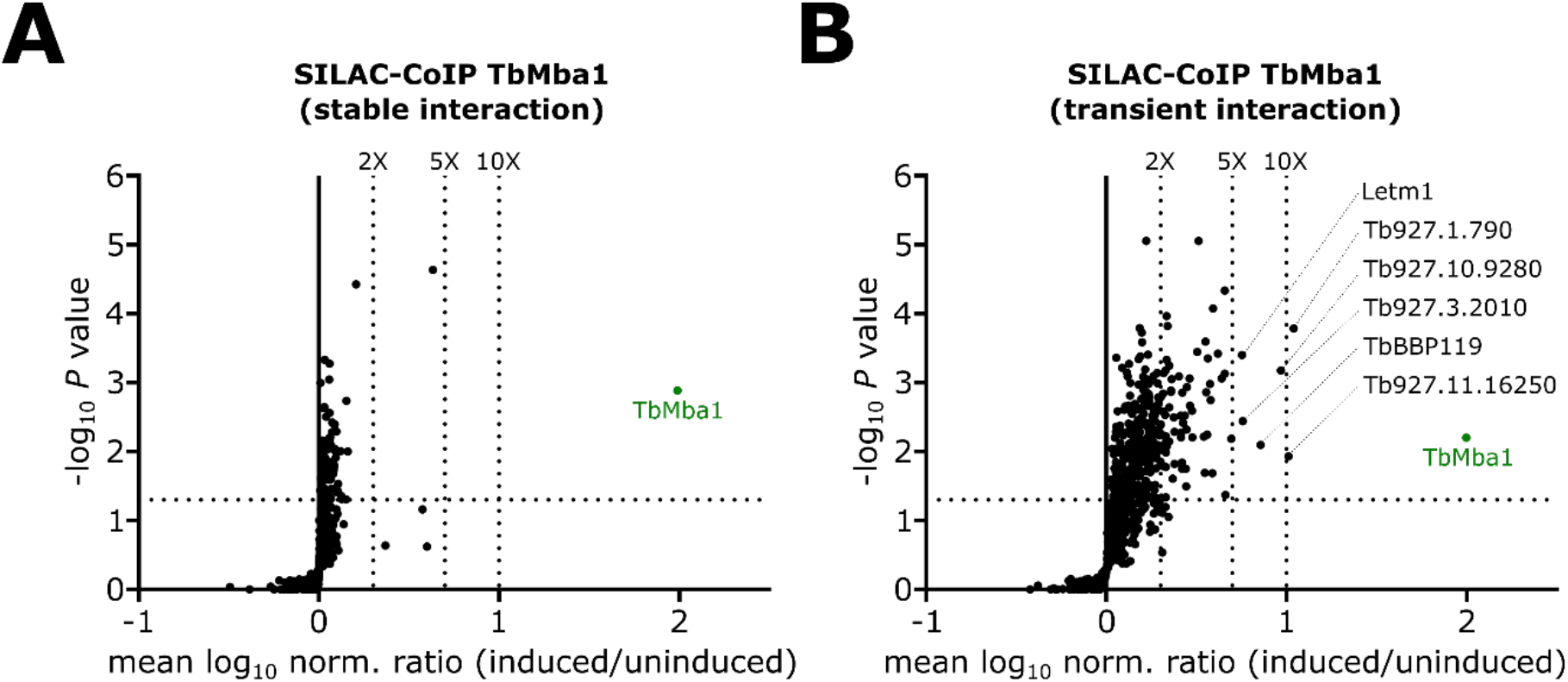
**A**: Volcano plot of data obtained by SILAC-based quantitative MS analysis of TbMba1-myc CoIPs, performed with mitochondria-enriched extracts. The induced and uninduced cells were mixed prior to performing the CoIP to detect stable interactions. The volcano plot shows the mean log^10^ of the normalized (norm.) ratio (induced/uninduced) of mitochondrial proteins that were detected in at least two of three independent biological replicates, plotted against the corresponding -log^10^ *P* values (one-sided t-test). TbMba1, the bait of the SILAC CoIP, was enriched 98-fold. The vertical dotted lines specify the indicated enrichment factors. The horizontal dotted line indicates a t-test significance level of 0.05. **B**: Volcano plot of SILAC-based quantitative MS analysis of TbMba1-myc CoIP as in A. Except, the CoIPs with induced and uninduced cells were performed separately and the eluates were only mixed prior to MS analysis to detect transient interactions. TbMba1 was enriched 100-fold. The vertical dotted lines specify the indicated enrichment factors. The horizontal dotted line indicates a t-test significance level of 0.05.

In order to detect possible transient interactions, the cultures of the TbMba1-myc exclusive expressor cell line were grown as described for the SILAC-CoIP above, but instead of mixing the heavy and light labelled cells prior to solubilization, the CoIPs were performed separately and only the resulting eluates were mixed^46^. The result is shown in the volcano plot in Fig. 5B, which depicts all mitochondrial proteins^43^ that were detected in at least two of three independent biological replicates. Again, the bait TbMba1 was 100-fold enriched (Table S3). Moreover, six proteins were enriched between five-and ten-fold. One of the enriched proteins was previously identified as TbLetm1 (Tb927.3.4920), the *T. brucei* ortholog of yeast Mdm38^47^. This is in line with the observation that in yeast, Mba1 and Mdm38 are interaction partners^19^ and suggests TbLetm1/Mdm38 might be involved in the biogenesis of mitochondrially encoded OXPHOS components. This is a novel function for TbLetm1 which was previously shown to be involved in K^+^/H^+^ homeostasis^23,47^. Intriguingly the latter function is also carried out by yeast Mdm38^48,49^. Thus, it is likely that TbLetm1 and yeast Mdm38 are functionally conserved. Another previously characterized protein that was enriched about ten-fold was the trypanosome-specific Tb927.11.16250^50^ which contains two pentatricopeptide repeats (PPR) (150-210 aa)^51^. PPR proteins are sequence-specific RNA-binding proteins that are involved in organellar gene expression in all eukaryotes^52^. In line with this, Tb927.11.16250 selectively stabilizes the mRNAs of mitochondrially encoded CoxI and CoxII^50^. Recent BLAST searches show that Tb927.11.16250 also contains a PilT N-terminus (PIN) domain (355-550 aa), which in many cases has endoribonuclease activity. Other enriched proteins include the trypanosomatid-specific mitochondrial proteins of unknown function Tb927.1.790, Tb927.10.9280 and Tb927.3.2010 (Fig. 5B). Moreover, the basal body protein BBP119 was also detected. However, BBP119 is a cytosolic protein and therefore likely a contaminant.

In summary, even though TbMba1 does not form a stable complex with other proteins, it transiently interacts with other proteins including TbLetm1 and CoxI/CoxII mRNA stabilisation factor Tb927.11.16250.

## Discussion

The IM is a particularly protein-rich membrane and contains the five protein complexes of the OXPHOS pathway responsible for ATP production, the most iconic function of mitochondria. The majority of the subunits of the OXPHOS complexes is encoded in the nucleus, synthesized in the cytosol and imported into mitochondria, like most other mitochondrial proteins. However, genes encoding for the enzymatic core subunits of the OXPHOS system are often retained in the mtDNA^1^ and the corresponding proteins are co-translationally inserted into the IM by membrane-associated mitoribosomes^10^. In *S. cerevisiae*, the interaction of the mitoribosomes with the IM is mediated by at least four factors: the insertase Oxa1^53^ via its C-terminus, the IM-associated Mba1^18^, and the two integral IM proteins Mdm38^20^ and Mrx15^54^. Moreover, the expansion segment 96-ES1 of the mitochondrial 21S rRNA has also been shown to form a contact site with the IM^21^. The four proteins mentioned above are widely conserved in a broad spectrum of eukaryotes, indicating that they might already have been present in the last eukaryotic common ancestor (LECA)^40^.

Here, we present a functional analysis of the trypanosomal Mba1 orthologue TbMba1. We show that TbMba1, as its yeast counterpart^16^, is a mitochondrially localized peripheral membrane protein. However, in contrast to yeast Mba1, whose absence resulted in a reduced growth rate on non-fermentable carbon sources only^16^, depletion of TbMba1 caused a strong reduction of growth of PCF *T. brucei* grown under standard condition in medium containing glucose^55^. This suggests that in PCF *T. brucei*, TbMba1 has a more essential role than yeast Mba1, possibly compensating for the absence of a Mrx15 orthologue which surprisingly has not been detected in Euglenozoa^40^, a subgroup of the Discoba which includes *T. brucei*^24^, even though it is found in essentially all other eukaryotic groups.

Depletion of TbMba1 causes reduced levels of many components of the OXPHOS system most of which are subunits of complex IV. The high sensitivity of complex IV to reduced TbMba1 levels can be explained by the fact that the *T. brucei* mitochondrial genome encodes three of its subunits (CoxI, II and III), whereas for the cytochrome bc1 complex, also termed complex III, and ATP synthase, also termed complex V, only a single subunit each is mitochondrially encoded. Interestingly, the steady state levels of MRPs remain unchanged in the absence of TbMba1 and so does the IM-association of the mitoribosomes. This suggests that the essential role of TbMba1 in PCF *T. brucei* is not connected to the integrity of the mitoribosomes but directly related to the co-translational integration of newly synthesized mitochondrially encoded proteins into the IM. This also indicates that while TbMba1 might contribute to the maintenance of a physical connection between mitoribosomes and the IM, this function is not essential.

Intriguingly, deleting both alleles of the TbMba1 gene does not affect growth of BSF trypanosomes. This makes sense since BSF *T. brucei* does not express respiratory complexes I to IV and therefore cannot perform OXPHOS. Instead, BSF cells produce their ATP via glycolysis^32,33^. Thus, no co-translational integration into the IM of mitochondrially encoded subunits of OXPHOS complexes is necessary. However, complex V is an exception, it is essential also in the BSF, not as an ATP synthase but as an enzyme that consumes ATP to maintain the membrane potential^38,39^. ATP synthase subunit a, also termed subunit 6, an essential component of the F_o_ moiety of the ATP synthase, is mitochondrially encoded and therefore must be inserted into the IM even in the absence of TbMba1. In yeast, IM insertion of ATPase subunit 6 is mainly mediated by Mdm38 in concert with Oxa1. Since orthologues of these two proteins are found in the *T. brucei* mitochondrion, the same may be the case for the trypanosomal ATP synthase subunit a.

The mitoribosomes of trypanosomes are very unusual. They have among the shortest known rRNAs of any translation system and 127 MRPs, a number that is much higher than that of any other ribosome^35,56–59^. Half of these proteins are specific for trypanosomes and its relatives. However, uL22 and uL29, the two MRPs that have been previously shown to crosslink to yeast Mba1^13,14,60^, are conserved in *T. brucei* and contribute to the shape of the exit tunnel region of the trypanosomal mitoribosome^35^. The same is the case for uL23 and uL24, which in yeast were shown to directly interact with the Oxa1 insertase^13,14,35,60^. It is possible that this region of the highly diverged trypanosomal mitoribosome is more restricted to evolutionary change because the association with the IM, which must be maintained for efficient co-translational IM integration, is based on conserved proteins like Mba1 and Oxa1.

Pulldown experiments using the cell line exclusively expressing tagged TbMba1 did not recover any proteins that were more than five-fold enriched suggesting that TbMba1 does not form a stable complex with other proteins. Thus, in agreement with the cryo-EM structure of the trypanosomal mitoribosome, TbMba1 itself is not a component of the mitoribosomes^35^. TbMba1 therefore behaves similarly to yeast Mba1 and differs from the mammalian Mba1 orthologue mL45, which is firmly integrated into the large mitoribosomal subunit^61^. TbMba1 however transiently interacts with five mitochondrial proteins, all of which were more than five-fold enriched in the postmix pulldown experiment. One of these proteins is TbLetm1, whose yeast orthologue Mdm38 was shown to function in the same processes as Mba1. The other previously characterized factor was Tb927.11.16250, a protein containing two PPR and a PIN domain, whose depletion specifically destabilizes CoxI and CoxII mRNAs^50^. This is in line with the observation that Mdm38 in yeast is involved in the recruitment of mRNA stabilization and translation factors^19^, and suggests that TbLetm1 might be involved in recruitment of Tb927.11.16250. Thus, Mba1, TbLetm1, Tb927.11.16250^50^ and possibly three other trypanosome-specific proteins of unknown function form a dynamic network that mediates and regulates co-translational insertion of OXPHOS complex subunits.

Thus, despite the highly diverged mitoribosome, the role of trypanosomal Mba1 and possibly Letm1 in mediating co-translational IM insertion of subunits of the OXPHOS system appears to be largely conserved between yeast and trypanosomes. It will be exciting to see how the more diverged RNA stabilization factors and translational regulators in different organisms fit into the dynamic interaction network defined by Mba1.

## Material and Methods

### Transgenic cell lines

Transgenic *T. brucei* cell lines are based on the procyclic form (PCF) strain 29-13 or the bloodstream form (BSF) strain New York single marker (NYsm)^62^. PCF *T. brucei* were grown in SDM-79 at 27°C in the presence of 10% (v/v) fetal calf serum (FCS)^55^. BSF *T. brucei* were grown in HMI-9 at 37°C in the presence of 10% (v/v) FCS and 5% CO_2_^63^.

RNAi cell lines were prepared using a modified pLew100 vector^62^, in which the phleomycin resistance gene was replaced by a blasticidin resistance gene. This vector allows the insertion of RNAi sequences in opposing directions with a 439 bp spacer fragment in between to form a stem loop. The TbMba1 (Tb927.7.4620) RNAis either target the nucleotides (relative to coding start) 322-756 (ORF) or 844-1094 (3’UTR). The TbMRPL22 (Tb927.7.2760) RNAi targets nucleotides (relative to coding start) 121-621 (ORF). The plasmids were linearized with NotI prior to stable transfection of PCF *T. brucei*. The TbMba1 dKO BSF cell line was generated by amplifying 500 nucleotides long sequences directly adjacent to the 5’ and 3’ ends of the coding sequence. The amplified 5’ and 3’ regions were ligated into pMOTag43M using XhoI/NdeI and BamHI/SacI, respectively, flanking the hygromycin resistance gene. The resulting plasmid was digested using XhoI/SacI prior to stable transfection of NYsm cells to replace the first TbMba1 allele with the hygromycin resistance gene. The second TbMba1 allele was replaced analogically by the blasticidin resistance gene using pBLUESCRIPT II KS+ instead of pMOTag43M.

To generate the TbMba1-myc exclusive expressor, first, a triple c-myc tagged variant of TbMba1 was produced (TbMba1-myc). The ORF of TbMba1, excluding the stop codon, was amplified by PCR and ligated into a modified pLew100 vector, containing a C-terminal triple c-myc tag and a stop codon. This plasmid was linearized with NotI prior to stable transfection into the 3’UTR RNAi cell line of TbMba1. Since the ectopically expressed TbMba1-myc does not contain the endogenous 3’UTR, it is resistant to the 3’UTR RNAi. The exclusive expressor cell line was induced with tetracycline two days before experiments to give the RNAi enough time to take effect. The in-situ HA-tagged mL78 (Tb927.10.11050) and L20 (Tb927.11.10170) variants (mL78-HA/L20-HA) were produced according to Oberholzer et al. (2006)^64^.

### RNA extraction and northern blotting

Total RNA was extracted from uninduced and two days induced RNAi cells, using acid guanidinium thiocyanate-phenol-chloroform extraction according to Chomczynski and Sacchi (1987)^65^. Equal amounts of RNA were separated on a 1% agarose gel containing 20 mM MOPS buffer and 0.5% formaldehyde. Gel-purified PCR products, corresponding to the respective RNAi inserts, were used as templates for the Prime-a-Gene labelling system (Promega) to radioactively label the northern probes.

### Antibodies

Commercially available primary antibodies used: mouse anti-c-myc (Invitrogen, dilution immunoblot (IB) 1:2’000, dilution IF 1:100), mouse anti-HA (Enzo Life Sciences AG, dilution IB 1:5’000) and mouse anti-EF1α (Merck Millipore, dilution IB 1:10’000). The polyclonal rabbit anti-ATOM40 (dilution IB 1:10’000, dilution IF 1:1’000), polyclonal rabbit anti-Cytochrome c (dilution IB 1:100), polyclonal rabbit anti-CoxIV (dilution WB 1:1’000), polyclonal rat anti-TbTim17 (dilution IB: 1:300), were previously produced in our laboratory^26,66^. The monoclonal mouse anti-TbmHsp70 (dilution IB 1:1,000) was kindly provided to us by Ken Stuart^67^. Commercially available secondary antibodies used: goat anti-mouse IRDye 680LT conjugated (LI-COR Biosciences, dilution IB 1:20’000), goat anti-Rabbit IRDye 800CW conjugated (LI-COR Biosciences, dilution IB 1:20’000), goat anti-mouse Alexa Fluor 596 (ThermoFisher Scientific, dilution IF 1:1’000), goat anti-rabbit Alexa Fluor 488 (ThermoFisher Scientific, dilution IF 1:1’000), HRP-coupled goat anti-mouse (Sigma, dilution IB 1:5’000). The BN-PAGE immunoblots were developed using the commercial kits SuperSignal West Pico PLUS or SuperSignal West Femto MAXIMUM.

### Immunofluorescence microscopy

2×10^6^ Cells were harvested, washed and resuspended in PBS, and then distributed on slides. They were allowed to settle on the plate for 10 min. After the incubation, they were fixed using 4% paraformaldehyde, and then permeabilized using 0.2% Triton X-100. The slides were incubated with the primary and secondary antibodies, described above, prior to picture acquisition using a DMI6000B microscope equipped with a DFC360 FX monochrome camera and LAS X software (Leica Microsystems).

### Digitonin extraction

A crude mitochondrial fraction was separated from a crude cytosolic fraction by incubating 1×10^8^ cells for 10 min on ice in 20 mM Tris-HCl pH 7.5, 0.6 M sorbitol, 2 mM EDTA containing 0.015% (w/v) digitonin. Centrifugation (5 min, 6’800 g, 4°C) separates a mitochondria-enriched pellet from the cytosol-containing supernatant. 2.5×10^6^ cell equivalents of each fraction were subjected to SDS-PAGE and immunoblotting to show mitochondrial localization of TbMba1-myc.

### Alkaline carbonate extraction

To separate soluble from membrane-integral proteins, a mitochondria-enriched pellet, obtained by digitonin extraction, was incubated for 10 min on ice in 100 mM Na_2_CO_3_ at pH 11.5 and centrifuged (100’000 g, 4°C, 10 min). Equal cell equivalents of each sample were analysed by SDS-PAGE and immunoblotting.

### Triton extraction

To test whether proteins were aggregated or not, the pellet from an alkaline carbonate extraction was incubated for 10 min on ice in 100 mM Na_2_CO_3_ at pH 11.5 containing 1% Triton X-100. After centrifugation (100’000 g, 4°C, 10 min), the supernatant and pellet were separated. Equivalents of each sample were analysed by SDS-PAGE and immunoblotting.

### SILAC RNAi

For the SILAC experiments, the cells were grown for four days in SDM-80^30^ containing 5.55 mM glucose, 10% dialyzed FCS (BioConcept, Switzerland), and either light (^12^C_6_/^14^N_χ_) or heavy (^13^C_6_/^15^N_χ_) isotopes of arginine (1.1 mM) and lysine (0.4 mM) (Euroisotop). Three biological replicates of the light/heavy pair were grown for each experiment. Either the light or heavy culture of each pair was induced with tetracycline three days before the harvest of TbMba1 RNAi cells or TbMRPL22 RNAi cells. 2□10^8^ cells of the heavy culture were mixed with 2×10^8^ cells of the light culture and a mitochondria-enriched pellet was obtained by digitonin extraction as described above. The triplicates were analyzed by liquid chromatography-mass spectrometry (LC-MS) as described below.

### SILAC CoIP

Six light/heavy pairs of the TbMba1-myc exclusive expressor cell line were grown for four days in SILAC medium as described above. Two days before harvest, either the light or heavy culture of each pair was induced with tetracycline. To identify stable interactions, three of the light/heavy pairs were prepared as follows: 2×10^8^ cells of the heavy culture were mixed with 2×10^8^ cells of the light culture and a mitochondria-enriched pellet was obtained by digitonin extraction as described above. The pellet was solubilized in lysis buffer (20 mM Tris-HCl, pH 7.4, 0.1 mM EDTA, 100 mM NaCl, 10% glycerol, 1% (w/v) digitonin, 1X Protease Inhibitor mix (EDTA-free, Roche)) for 15 min on ice. After centrifugation (20,000 g, 4°C, 15 min), the supernatant was incubated with anti-myc affinity matrix (Roche) for 2 h at 4°C. After incubation, the supernatant was removed, and the resin-bound proteins were eluted by boiling the resin for 5 min in SDS loading buffer (2% SDS (w/v), 62.5 mM Tris-HCl pH 6.8, 7.5% glycerol (v/v), bromophenol blue). The proteins were precipitated using methanol and chloroform and resuspended again in loading buffer. The samples were run 1 cm into a 4-12% NuPAGE BisTris gradient gel (Life Technologies). The proteins were stained using colloidal Coomassie Brilliant Blue and the gel piece containing the proteins was cut out. The proteins in the gel fragments were reduced using 5 mM Bond-Breaker TCEP solution (Thermo Scientific, Product No. 77720), alkylated using 100 mM iodacetamide (Sigma-Aldrich, Product No. I6125), and digested in-gel with trypsin. The triplicates were then analysed by LC-MS as described below.

To identify transient interactions^46^, the same procedure as above was used, but instead of mixing the cells in the beginning, the eluates of each light/heavy pair were mixed after elution from the anti-myc affinity matrix.

### LC-MS and data analysis

Mitochondria-enriched fractions obtained from SILAC-labeled tetracycline-induced and -uninduced TbMba1 RNAi or TbMRPL22 RNAi cells were prepared for LC-MS analysis as described before^68^. In brief, mitochondrial fractions were resuspended in urea buffer (8 M urea/50 mM NH_4_HCO_3_), followed by reduction and alkylation of cysteine residues, dilution of samples to a final urea concentration of 1.6 M using 50 mM NH_4_HCO_3_, and tryptic digestion. Peptide mixtures were desalted using StageTips, dried *in vacuo* and reconstituted in 0.1% (v/v) trifluoroacetic acid. Sample were analyzed in technical duplicates by nano-HPLC-ESI-MS/MS on an Orbitrap Elite (Thermo Fisher Scientific) coupled to an UltiMate 3000 RSLCnano HPLC system (Thermo Fisher Scientific), which was equipped with PepMap C18 precolumns (Thermo Scientific; length, 5 mm; inner diameter, 0.3 mm; flow rate, 30 μl/min) and an Acclaim PepMap C18 reversed-phase nano-LC column (Thermo Scientific; length, 500 mm; inner diameter, 75 μm; particle size, 2 μm; pore size, 100 Å; flow rate, 0.25 μl/min). Peptides were separated using a binary solvent system consisting of 4% (v/v) dimethyl sulfoxide (DMSO)/0.1% (v/v) formic acid (FA) (solvent A) and 48% (v/v) methanol/30% (v/v) acetonitrile (ACN)/4% (v/v) DMSO/0.1% (v/v) FA (solvent B). A gradient ranging from 1% - 25% solvent B in 115 min followed by 25% - 45% B in 110 min, 45% - 60% B in 50 min, 60% - 80% B in 20 min, 80% - 99% B in 10 min and 5 min at 99% B was applied for peptide elution. Peptide mixtures generated in TbMba1-myc SILAC CoIP experiments were analyzed by LC-MS using the same instrument setting as described above except that the RSLCnano HPLC was equipped with nanoEase M/Z Symmetry C18 precolumns (Waters; length, 20 mm; inner diameter, 0.18 mm; flow rate, 10 μl/min) and a nanoEase M/Z HSS C18 T3 column (Waters; length, 250 mm; inner diameter, 75 μm; particle size, 1.8 μm; packing density, 100 Å; flow rate, 300 nl/min). Peptides were eluted using 0.1% (v/v) FA (solvent A) and 50% (v/v) methanol/30% (v/v) ACN/0.1% (v/v) FA (solvent B) as solvent system and the following gradient: 7% - 55% solvent B in 175 min followed by 55% - 95% B in 35 min and 5 min at 95% B. Mass spectrometric data were acquired in data-dependent mode, applying the following parameters: mass range of *m/z* 370 to 1,700, resolution of 120,000 (at *m/z* 400), target value of 10^6^, maximum injection time of 200 ms for MS survey scans. Up to 25 of the most intense precursor ions (z ≥ +2) were selected for further fragmentation by low energy collision-induced dissociation in the linear ion trap with a normalized collision energy of 35%, an activation q of 0.25, an activation time of 10 ms, a target value of 5,000, a maximum injection time of 150 ms, and a dynamic exclusion time of 45 s. For protein identification and SILAC-based quantification, MaxQuant/Andromeda^69,70^ (versions 1.5.3.12, 1.5.4.0, and 1.6.5.0 for data obtained in TbMba1 RNAi, TbMRPL22 RNAi, and TbMba1-myc CoIP experiments, respectively) and the fasta file for *T. brucei* TREU927 (downloaded from the TriTryp database; version 8.1) were used. Data were processed using MaxQuant default settings except that protein identification and quantification were based on ≥ 1 unique peptide and ≥ 1 ratio count, respectively. Lys0/Arg0 were set as light and Lys8/Arg10 as heavy labels. The mean of log^10^- transformed normalized protein abundance ratios were determined for proteins quantified in ≥ 2 independent biological replicates per dataset and a two-sided (SILAC RNAi experiments) or one-sided (TbMba1-myc CoIP experiments) Student’s t-test was performed. Information about the proteins identified and quantified are provided in Tables S1 - S3.

### Blue native-PAGE

To analyse native complexes, a mitochondria-enriched pellet obtained by digitonin extraction was used as starting material. This pellet was solubilized in solubilization buffer (20 mM Tris-HCl pH 7.4, 50 mM NaCl, 10% glycerol, 0.1 mM EDTA) containing 1% (w/v) digitonin and incubated on ice for 15 min. After centrifugation (20’000 g, 4°C, 15 min), the supernatant was separated on 4-13% gels. Before western blotting, the gel was incubated in SDS-PAGE running buffer (25 mM Tris, 1 mM EDTA, 190 mM glycine, 0,05% (w/v) SDS) to facilitate transfer of proteins.

### IM association of mitoribosomes

To separate a mitochondrial matrix fraction from a mitochondrial membrane fraction, 1.5×10^8^ TbMba1 ORF RNAi cells that also in-situ express either mL78-HA or L20-HA were induced for up to four days with tetracycline. Mitochondria-enriched pellets were obtained using digitonin extraction as described above. The pellets were resuspended in 150 μl 10 mM MgCl_2_ and then flash frozen in liquid nitrogen and thawed at 25°C ten times. After centrifugation (10’000 g, 5 min, 4°C), the supernatant was separated from the pellet, which was resuspended in 150 μl 10 mM MgCl_2_. 37.5 μl 5X loading buffer (10% SDS (w/v), 312.5 mM Tris-HCl pH 6.8, 12.5% β-mercaptoethanol (v/v), 37.5% glycerol (v/v), bromophenol blue) was added to both fractions and incubated 5 min at 95°C. 15 μl of both fractions were analysed by SDS PAGE and immunoblotting.

## Acknowledgements

We thank Bettina Knapp for technical assistance. Work in the lab of B.W. was supported by Germany’s Excellence Strategy (CIBSS – EXC-2189 – Project ID 390939984). Work in the lab of A.S. was supported in part by NCCR RNA & Disease, a National Centre of Competence in Research (grant number 205601) and by project grant SNF 205200 both funded by the Swiss National Science Foundation.

## Supplementary figures and legends

**Figure S1.**
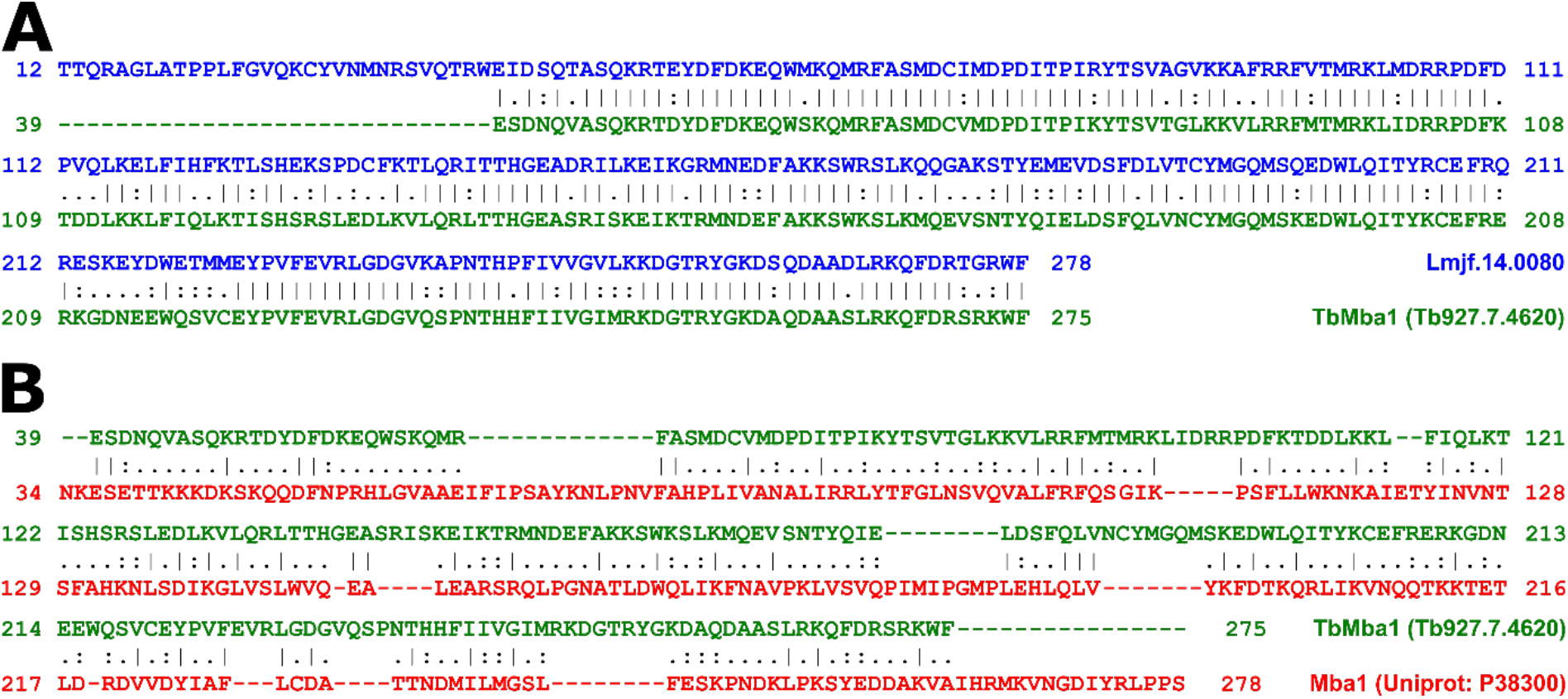
**A**: Alignment of Lmjf.14.0080 (blue) and TbMba1 (green), using EMBOSS Needle with BLOSUM62 as substitution matrix^71^. The N-terminal presequences, as predicted by MitoProt II server (https://ihg.gsf.de/ihg/mitoprot.html), were excluded (Lmjf.14.0080: amino acids (aa) 1-11, TbMba1: aa 1-38). (|) identical residue, (.) conserved change, (:) part of residue is similar but not that conserved^72^. **B**: Alignment as in A but between TbMba1 (green) and yeast Mba1 (Uniprot: P38300, red). The predicted N-terminal presequences were again excluded (Mba1: aa 1-33).

**Figure S2.**
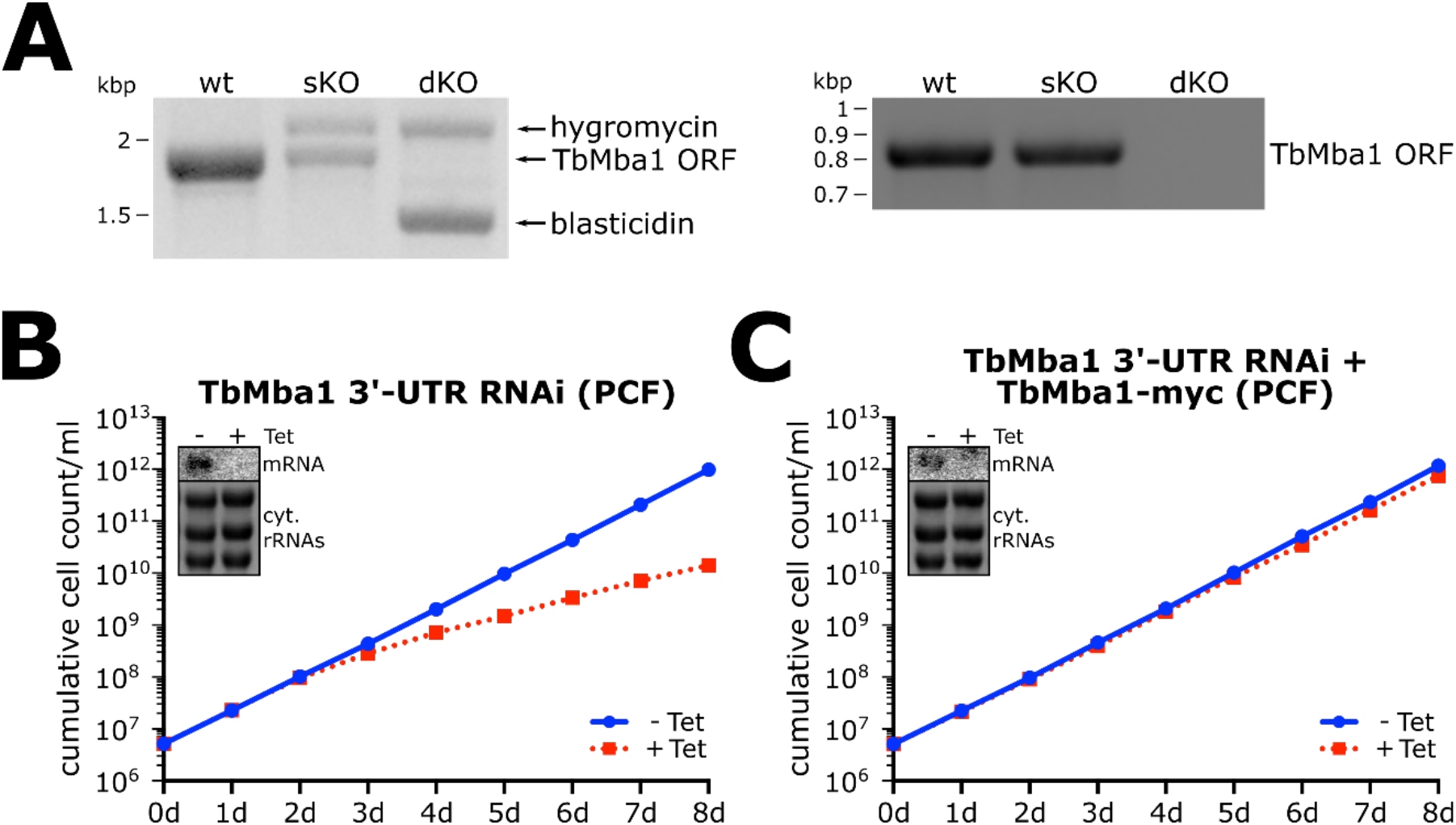
**A**: Left panel: confirmation of the TbMba1 single knockout (sKO) and double knockout (dKO) NYsm BSF cell lines. The used primers start 550 bp upstream and downstream of the replaced allele. The first TbMba1 allele was replaced with the hygromycin resistance cassette, the second allele with the blasticidin resistance cassette. Right panel: PCR using primers within the ORF to exclude the possibility that the ORF of TbMba1 is present in another locus. **B**: Growth curve of uninduced (-Tet) and induced (+Tet) TbMba1 RNAi cell line targeting the 3’-UTR of the mRNA coding for TbMba1. Inset show northern blot of total RNA from uninduced and two days induced cells, probed for the TbMba1 ORF. Ethidium bromide (EtBr)-stained cytosolic (cyt.) rRNAs serve as loading control. **C**: Growth curve of the uninduced and induced TbMba1-myc exclusive expressor cell line. The 3’-UTR RNAi cell line from B ectopically expresses TbMba1-myc at the same time. Insets as in left panel.

**Figure S3.**
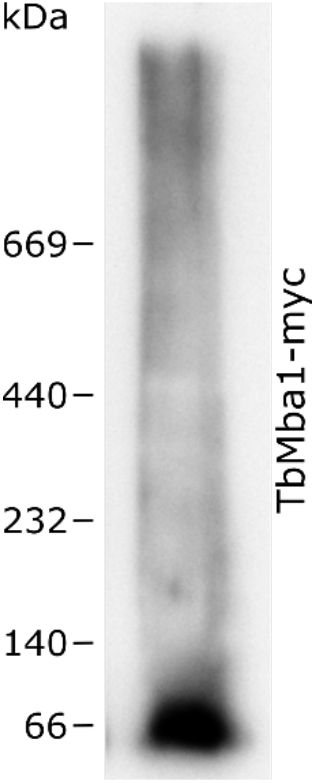
BN-PAGE and immunoblot analysis of a solubilized mitochondrial fraction of the TbMba1-myc exclusive expressor cell line, induced for two days. The immunoblot was probed with anti-myc antibody.

